# Neglected predatory insects trigger potential Key Biodiversity Areas in threatened coastal habitats

**DOI:** 10.1101/2024.12.02.626341

**Authors:** Aleida Ascenzi, Dario Nania, Andrea Cristiano, Davide Badano, Michela Pacifici, Pierfilippo Cerretti

**Author notes:** Corresponding authors Aleida Ascenzi Dario Nania.

## Abstract

Key Biodiversity Areas (KBAs) are poised to become a powerful tool for identifying regions that host unique biodiversity. With their great diversity, insects hold significant potential as indicators for global KBA mapping, even in highly specialized and narrowly distributed habitats. For instance, species adapted to fragmented ecosystems like coastal sand dunes—among the most heavily impacted habitats worldwide—can serve as critical indicators to trigger KBAs in these fragile environments. Despite their relevance as indicators, the inclusion of insects in KBA assessments remains limited, especially for neglected insect species such as antlions. We tested selected KBA criteria on 26 antlion and owlfly species (Neuroptera: Myrmeleontidae) in Italy, including dunes specialists, and performed COI based genetic analysis to identify potential weaknesses of the assessment. Several endemic and dune specialist species trigger potential KBAs, showing limited (< 20% of their extent) overlap with the current protected area network, confirming the great value of these taxa in narrowly distributed habitats. Genetic data provides evidence of misidentification for widely distributed species. We advise for the integration of both spatial and genetic data to increase reliability of potential KBA assessments using neglected insect taxa.

## 1. INTRODUCTION

Key Biodiversity Areas (KBAs) *are sites contributing significantly to the global persistence of biodiversity* (IUCN 2016), recognized as a global standard for identifying relevant areas for site-based conservation (Chen & Wu 2019). KBAs have been already integrated into global conservation policies, for instance by serving as indicators for selected targets for the achievement of the Sustainable Development Goals by 2030 (CBD 2022). Despite their widespread adoption, KBAs exhibit a significant taxonomic bias, disproportionately focusing on plants and birds (BirdLife International 2021), while largely overlooking critical insect biodiversity, with only 6.04% of existing KBAs are triggered by invertebrate species (Chowdhury et al. 2022). This underrepresentation is particularly concerning as invertebrates, including insects, comprise the majority of biodiversity and are essential for ecosystem functioning (Reiss et al. 2009; McCary et al. 2021). Many insect taxa are experiencing dramatic declines, which could lead to the extinction of 40% of the world’s insect species in the coming decades (Sánchez-Bayo & Wyckhuys 2019).

Ensuring genetic and phylogenetic diversity is essential for the persistence of threatened species’ populations (Sgrò et al. 2011; Brooks et al. 2015), and integrating genetic and distributional data is fundamental for understanding biodiversity patterns and processes at multiple scales (Soltis & Soltis 2016; Arribas et al. 2021). KBA methodology includes ‘distinct genetic diversity’ as a metric under Criteria A1, B1 and B2 (KBA Standards and Appeals Committee of IUCN SSC/WCPA 2022). This criterion allows the identification of KBAs in cases where a site contains a significant proportion of a species’ global genetic diversity, even when the population size within the site does not meet other thresholds. However, this approach is rarely applied in practice for identifying potential KBAs, particularly for insect taxa. The underutilization of genetic criteria exacerbates the challenges posed by the lack of comprehensive genetic and distributional data for many arthropod groups.

DNA barcoding has emerged as a pivotal tool for studying arthropod diversity, allowing species identification (Hebert et al. 2003; Hebert 2016) and overcoming the limitations of traditional methods based on morphology (Behrens-Chapuis et al. 2021). However, despite its potential to significantly contribute to biogeographical analyses of insect taxa (Greenstone 2011; da Silva et al. 2023), major challenges remain. These include undersampling of arthropod taxa (Liu et al. 2022; Souther et al. 2024), limited access to private data (Willemse et al. 2019), geographic and temporal biases (Cardoso & Leather 2019; Rocha-Ortega et al. 2021; Sánchez-Fernández et al. 2021), and taxonomic impediments (Engel et al. 2021; Vinarski 2020). Such limitations hinder the availability of comprehensive distributional data for insect taxa (Rocha-Ortega et al. 2021), thereby limiting our understanding of their geographic and ecological ranges. These barriers can undermine the accuracy of conservation assessments that rely on robust population estimates or precise delineations of species’ global distributions, such as those employed in the KBA framework (IUCN, 2016). As a result, these assessments often rely on higher taxonomic levels instead of species-specific units (Heino & Soininen 2007), leading to the deprioritization of insects in conservation planning and management. A first assessment to identify potential KBAs for insect taxa was conducted by Nania et al. (2024a) for bumblebees (Hymenoptera: Apidae: *Bombus*), who highlighted major challenges when applying KBA criteria to insect species.

Myrmeleontidae (Neuroptera) are a rarely investigated group in conservation studies, and there are still important gaps in our knowledge of their taxonomy and distribution. Myrmeleontidae can be divided in two functional groups: i) antlions, including Myrmeleontinae, Dendroleontinae, Nemoleontinae, Ascalaphinae Dimarini, and Palparini, characterized by weak flying adults and largely fossorial larvae; and ii) owlflies, comprising Ascalaphinae Ululodini, Haplogleniini and Ascalaphini known for their strong flying adults and larvae with camouflaging adaptations (Badano et al. 2018; Machado et al. 2019). Some species have specialized larval niche requirements (Stange & Miller, 1990; Stange et al. 2003), leading to restricted distributions, while others are more generalist and exhibit wide distribution ranges (Mansell & Erasmus 2002; Hévin et al. 2023; Zheng et al. 2024). Myrmeleontidae are a significant component of insect fauna in arid environments, with most species closely associated with sandy habitats, including coastal dunes (Miller & Stange 1985; Mansell 1999). These habitats are increasingly threatened by degradation, fragmentation, and climate change (Mentaschi et al. 2018).

Coastal sand dunes are among the most impacted habitat types globally (Brown & McLachlan 2002; Schlacher et al. 2007; EEA 2009). According to the European Commission Directorate-General for Environment (2016), 45% of coastal habitats of Europe are threatened, and in the Mediterranean region, 75% of all dune systems have been destroyed due to extensive tourism development (German Federal Agency for Nature Conservation) and several ecosystems have been assessed as endangered (https://www.assessments.iucnrle.org). In Italy, 88% of dune habitats are in poor conservation status (Prisco et al. 2020). At the national level, the Red List of Ecosystems classifies herbaceous, shrubby, and halohygrophilous coastal ecosystems as Vulnerable, whereas psammophilous ecosystems are considered Critically Endangered on the mainland and Endangered on the islands (Blasi et al. 2023). Despite this, most beaches are not under any marine or terrestrial management initiatives.

The association of Myrmeleontidae with threatened coastal habitats, their varying distribution range, and unresolved taxonomy make them valuable candidates for applying the KBA methodology. In this study, we (i) apply selected KBA criteria (B1, B2 and B3) to 26 species of antlions and owlflies to identify potential KBAs in Italy; (ii) estimate the extent of overlap between these potential KBAs and the Italian protected areas (PAs), Natura 2000 sites, and the current map of Italian KBAs; and (iii) investigate potential species-level misidentifications that could impact the identification of KBAs by analysing barcode gaps in geotagged cytochrome oxidase I (COI-5P) sequences available from the Barcode of Life Data System (BOLD; Ratnasingham 2007) and NCBI (National Center for Biotechnology Information) databases.

## 2. MATERIALS AND METHODS

### 2.1 Distributional data

The species included in the study were selected from the Italian checklist of Myrmeleontidae and Ascalaphidae, focusing on Euro-Mediterranean species (Online Resource 1, Table S1). We built a comprehensive dataset of occurrence points for 7 owlflies and 19 antlion species to delineate their global distributional ranges. We gathered distribution data from the literature, private collection specimens and image-validated records from GBIF (Global Biodiversity Information Facility) (the reference list of occurrences gathered from the literature is available at Online Resource 1, Section S3). Since the potential KBA assessment was performed in Italy, we refined the dataset for the Italian territory by integrating research-grade occurrences from iNaturalist, including only image-validated records (www.inaturalist.org). We georeferenced the maps of Neuroptera distribution provided by Aspöck et al. (1980) using the Georeferencer extension in QGIS (QGIS.org 2022), with a polynomial 3 transformation type and nearest neighbour resampling. From the georeferenced maps we then extracted occurrence points of each species, approximating a spatial accuracy of 10 kilometres (i.e. the georeference points were identified within a circular area of approximately 10 kilometres of radius). For recently described species (i.e., *Myrmeleon mariaemathildae* Pantaleoni, Cesaroni & Nicoli Aldini, 2010 and *My. punicanus* Pantaleoni & Badano, 2012) we only considered literature and GBIF records. The records span from 1800 to 2023. We acknowledge that some of the old records might reflect environmental conditions that are substantially different from the current ones, directly affecting the probability of recording species’ presences. These data were used to estimate the limits in the spatial and climatic distribution of the various species. We here assume that the true species distribution may vary to a relatively negligible extent and considered three different approaches that can be used as proxies to estimate the species’ global population size for scoping KBAs (IUCN 2016): geographic distribution ranges, Area of Occupancy (AOO) and Area of Habitat (AOH).

### 2.2 Global population size estimate

We produced a set of maps representing the species’ global distribution ranges using a regular 10 x 10 km cell grid. For each species, we extracted the grid cells that overlapped with the occurrence points. The global distribution range is represented by all retained cells for each species. This approach has been adopted extensively to map species distributions based on occurrence points (Hickling et al. 2005; Nania et al. 2022).

We produced AOO maps following the Mapping Standards and Data Quality for the IUCN Red List Categories and Criteria (IUCN 2021). The Area of occupancy of a species is a scaled metric used to map the habitat occupied by a species (IUCN 2021). We overlapped a regular 2 x 2 km cell grid over the species occurrence points and extracted only grid cells having at least one species occurrence falling within them. The total surface area of the selected cells is the AOO of the species and corresponds to an estimate of the extent of habitat occupied by the species (IUCN Standards and Petitions Committee 2024).

A set of AOH maps was obtained by combining distribution ranges with a land cover map and altitude data (Brooks et al. 2019; Rondinini et al. 2011). The same procedure was previously adopted to produce AOH maps for bumblebees in Italy (Nania et al. 2024a). Land-cover categories of the CGLS-LC100 Copernicus Land Cover Map (Buchhorn et al. 2020) were used as habitat surrogates. We estimated altitudinal minimum and maximum limits for the species based on literature (Steffan 1975; Aspöck et al. 1980; Gepp 2010; Krivokhatsky 2011; Monserrat & Acevedo 2013; Badano & Pantaleoni 2014a,b). A table showing land cover categories used as proxies for the species’ habitat is available in the Online Resource 2, while a detailed description of how AOH maps were produced is available in the Online Resource 1 (Section S2). Although we produced maps with all three approaches, the subsequent steps were followed only for the distribution range maps due to unreliability of AOH and AOO maps (see Discussion section).

### 2.4 Potential KBAs

We selected three KBA criteria that can be evaluated by employing distribution range maps as proxy of the global population size of the species (IUCN 2016). The selected criteria were B1 (individual geographically restricted species), B2 (co-occurring geographically restricted species), and B3 (geographically restricted assemblages) (IUCN 2016; KBA Standards and Appeals Committee of IUCN SSC/WCPA, 2022). Criterion B1 refers to species with a significant proportion (i.e., ≥ 10%) of their global population restricted to a specific site. Criterion B2 concerns geographically restricted species, and it can be activated when a site contains ≥ 1% of the global population of at least two geographically restricted species. Criterion B3 can be triggered if a site hosts ≥ 0.5% of the global population of at least 5 species restricted to a specific ecoregion (IUCN 2016; KBA Standards and Appeals Committee of IUCN SSC/WCPA 2022).

To identify potential KBAs in Italy for our selection of species, we employed a systematic application of KBA criteria following Nania et al. (2024b). The method uses a moving grid with adjustable cell sizes to scope potential KBAs within the study area based on the species distribution data in the form of Area of habitat (AOH), Area of occupancy or distribution range. This is considered a first step in the KBA identification process, the second step is the KBA delineation and requires confirmation of the species’ presence in the site with a sufficient number of mature individuals (KBA Standards and Appeals Committee of IUCN SSC/WCPA 2022). We used a 10 x 10 km cell size grid, which has been previously used in KBA assessments for insects (Nania et al., 2024a). However, we also tested a 5 km x 5 km cell size grid to compare the output maps and test the stability of potential KBAs in terms of number of species regardless of the cell size. We identified potential KBAs using 10 x 10 kilometres occurrence cells as an estimate of species’ ranges and estimated the extent of overlap between the potential KBAs and national protected areas (NNB 2021), Natura 2000 sites map (EEA 2021), and the current KBA network in Italy (BirdLife International 2021).

All figures were created by assembling images derived from maps visualized in QGIS (v. 3.22.5), using a licensed version of Adobe Photoshop 2021.

### 2.5 Intraspecific diversity analyses

We downloaded 160 COI sequences for analysis, including 38 sequences from Ascalaphinae and 122 sequences from Myrmeleontinae, Dendroleontinae, and Nemoleontinae, from the BOLD and NCBInt databases. Sequences of the 5’ region of cytochrome c oxidase subunit I (COI-5P) were selected, aligned using MAFFT v7 (Katoh & Standley 2013), and trimmed to 497 bp (owlflies) and 316 bp (antlions), based on the minimum shared nucleotide positions within each subgroup. Barcodes from South Africa were excluded *a priori* due to inconsistencies with the known distribution ranges of the taxa. Fasta format file of the downloaded sequences is available in Online Resource 3. The remaining sequences were clustered into putative species using an uncorrected p-distance threshold of 3% (Ratnasingham & Hebert 2013), implemented via a Python script utilizing the Objective Clustering algorithm (Meier et al. 2006). Barcode gaps for Ascalaphinae and for Myrmeleontinae, Dendroleontinae, and Nemoleontinae, were assessed to investigate intraspecific and interspecific diversity, focusing on potential misidentifications and cryptic diversity. The results were visualized using two dendrograms for the two subgroups, and, when necessary, species-specific dendrograms.

## 3. RESULTS

### 3.1 Distributional data

Our dataset comprised 3,054 occurrence records, including 2,443 digitized from the literature and 162 from private collections (Online Resource 1, Table S1). For antlions, we also incorporated 326 occurrences from iNaturalist and 123 from GBIF. We generated global maps of AOO, AOH and geographic distribution ranges for the 26 species analyzed. The AOH maps demonstrated poor performance in validation tests (Online Resource 1, Section S2) and did not outperform a random distribution model. Using a 10 km x 10 km cell grid, we reconstructed 26 global distribution range maps, all of which are available in Online Resource 4.

### 3.2 Potential KBAs

Using 10 km x 10 km cell sized grids, sixteen species of lacewings (twelve antlions and four owlflies) triggered two of the three selected criteria (B1 and B2) (Table 1).

**Table 1.** List of the species of Myrmeleontidae studied, with details on triggered KBA criteria and intraspecific diversity.

| Species | Triggered KBA<br>criteria | Number of geotagged<br>COI-5P sequences<br>analysed | Maximum<br>intraspecific p-distance<br>(%) |
| --- | --- | --- | --- |
| **Ascalaphinae** |  |  |  |
| *Deleproctophylla australis* | B2 |  |  |
| *Libelloides coccajus* |  | 5 | 0.4 |
| *Libelloides corsicus* | B2 |  |  |
| *Libelloides lacteus* |  |  |  |
| *Libelloides latinus* | B2 |  |  |
| *Libelloides longicornis* |  | 30 | 1.41 |
| *Libelloides siculus* | B2 |  |  |
| *Palpares libelluloides* |  | 3 | 9.48* |
| **Myrmeleontinae** |  |  |  |
| *Myrmeleon gerlindae* | B2 | 6 | 1.9 |
| *Myrmeleon mariaemathildae* | B1, B2 | 16 | 0.32 |
| *Myrmeleon punicanus* | B1, B2 | 6 | 0.95 |
| *Solter liber* | B1, B2 | 5 | 0.32 |
| *Synclisis baetica* | B2 |  |  |
| Dendroleontinae |  |  |  |
| *Dendroleon pantherinus* |  | 3 | 12.03* |
| Nemoleontinae |  |  |  |
| *Creoleon corsicus* | B1, B2 |  |  |
| *Creoleon lugdunensis* |  | 14 | 0.63 |
| *Distoleon annulatus* |  | 9 | 0.63 |
| *Distoleon tetragrammicus* |  | 18 | 6.37* |
| *Macronemurus appendiculatus* |  | 12 | 1.27 |
| *Megistopus flavicornis* |  | 11 | 4.46* |
| *Megistopus lucasi* | B1, B2 |  |  |
| *Neuroleon arenarius* | B2 | 3 | 0.32 |
| *Neuroleon egenus* | B2 | 8 | 14.42* |
| *Neuroleon microstenus* | B2 |  |  |
| *Neuroleon nemausiensis* | B2 | 11 | 16.29* |
| *Neuroleon ochreatus* | B1, B2 |  |  |

\*Species exceeding the delimitation threshold (> 3% p-distance)

Specifically, potential KBAs were identified under criterion B2 for *Creoleon corsicus*, *Deleproctophylla australis*, *Libelloides corsicus*, *L. latinus*, *L. siculus*, *Megistopus lucasi*, *Myrmeleon gerlindae, My. mariaemathildae*, *My. punicanus* Pantaleoni & Badano, 2012*, Neuroleon arenarius*, *N. egenus*, *N. microstenus*, *N. nemausiensis*, *N. ochreatus*, *Solter liber*, and *Synclisis baetica*. *Creoleon corsicus*, *Me. lucasi*, *My. mariaemathildae*, *My. punicanus*, *N. ochreatus*, and *So. liber*, triggered potential KBAs under both criteria B1 and B2. Overall, the potential KBAs identified for myrmeleontids represent 1.16% of the Italian territory (3,484 km²) (Fig. 1).

**Fig. 1.**
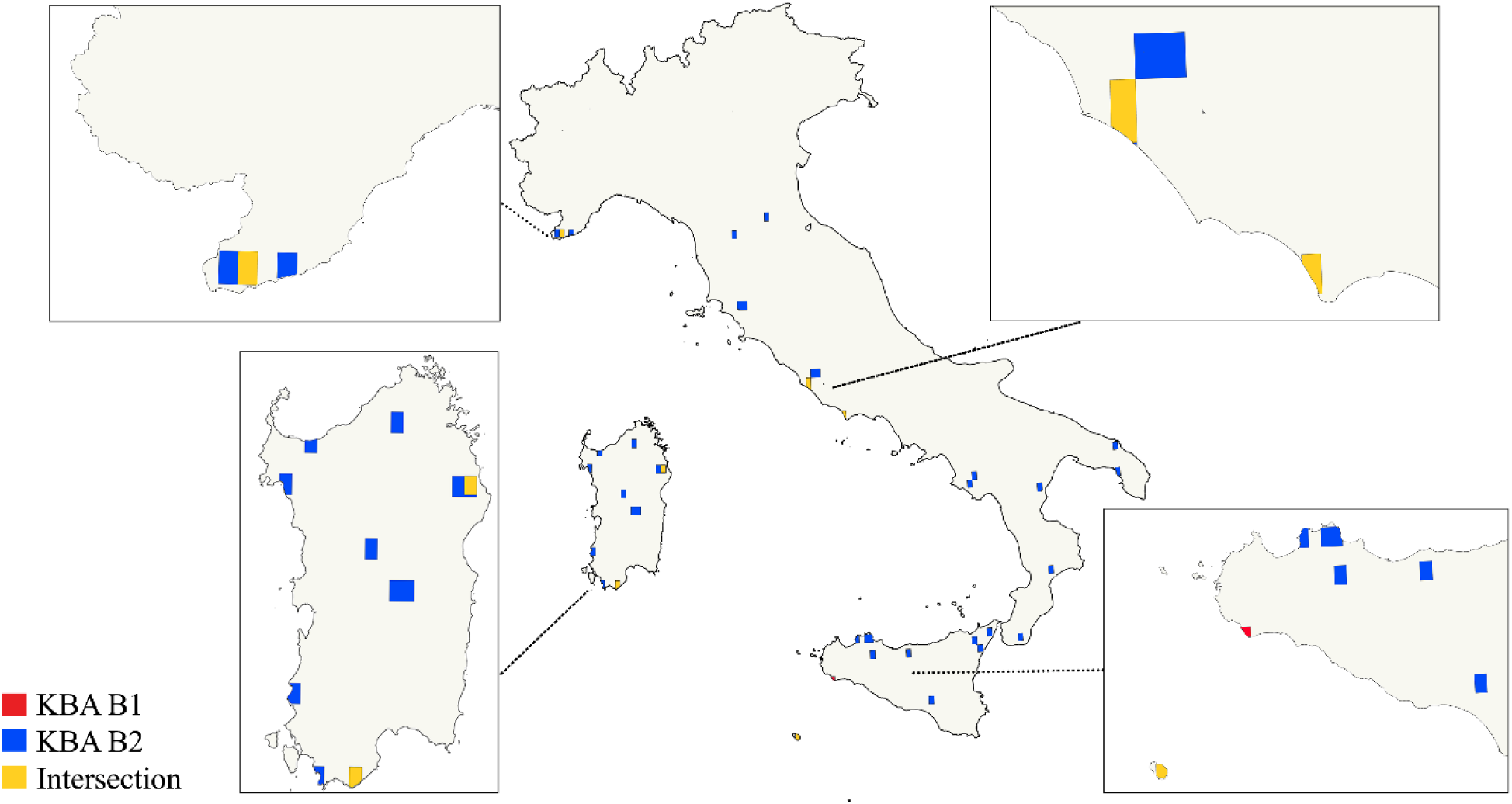
Potential KBAs in Italy for antlions and owlflies with details on triggered criteria

Potential KBAs were identified in north-west, central, and southern Italy, as well as on the islands of Sardinia, Sicily and Pantelleria. Many of these areas were associated with coastal dune habitats, particularly in the central Tyrrhenian region and in northern, southern, and south-western parts of Sardinia. Here, four dune-specialist species–*Myrmeleon mariaemathildae*, *Megistopus lucasi*, *Neuroleon ochreatus*, and *Synclisis baetica–*triggered potential KBAs (Fig. 2).

**Fig. 2.**
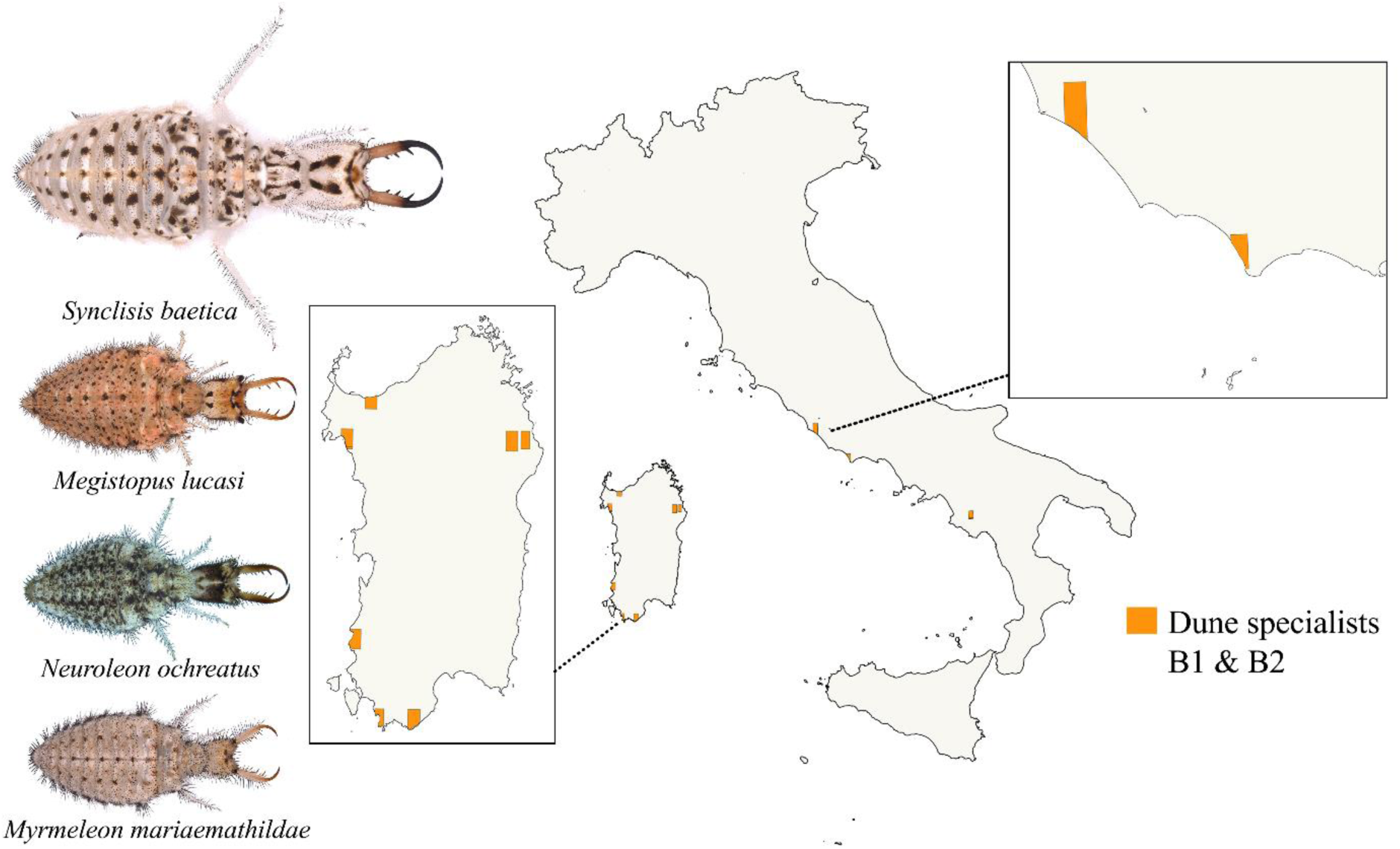
Potential KBAs in Italy for dune-specialist antlion species and their larval morphology

The identified potential KBAs overlapped with national protected areas for 18.67% of their extent, within Natura 2000 sites for 27.63%, and with current KBA maps for 19.94%. Maps of these nested areas are provided at Online Resource 1, Fig. S1, S2 and S3. Using 5 km x 5 km grid cells, four species triggered potential KBAs: *Myrmeleon punicanus* under criterion B1 and *My. punicanus, My. gerlindae*, and *Megistopus lucasi*, under criterion B2 (Online Resource 1, Fig. S4). The total area of potential KBAs was 261.52 km², representing 0.09% of the Italian territory. No KBAs were identified using criterion B3 at any resolution, as no area had at least 5 ecoregionally restricted species co-occurring.

The identified KBAs are concentrated in relatively well-preserved dune systems, though many are under strong anthropogenic pressure. Sampling biases may also influence these results, as extensively sampled sites correspond to identified KBAs.

All identified potential KBA maps are available at Online Resource 5.

### 3.3 Intraspecific diversity analysis

A total of 160 COI sequences were analysed, yielding 4 mOTUs for Ascalaphinae and 21 for Myrmeleontinae, Dendroleontinae and Nemoleontinae at a 3% p-distance threshold (Online Resource 1, Fig. S5 and S6). Barcode gaps analysis revealed varying levels of intracluster diversity across nominal species (Table 1), with some species showing high (putative) intraspecific divergence, suggesting cryptic diversity or misidentifications. For antlions (*sensu latu*), several species exhibited significant intraspecific divergence:

*Distoleon tetragrammicus*: specimens from Italy, Germany, and Portugal showed a p-distance of 6.4% from those in Slovakia and Azerbaijan. One specimen from Austria (MG334611.1) clustered with the eastern-most specimens, but additional sequences are needed to confirm this placement;

*Megistopus flavicornis*: A Russian specimen (OR591047.1) showed a 4.5% p-distance from specimens from Portugal;

*Distoleon pantherinus*: Of the three available sequences, those from Slovenia and Austria showed no genetic differences, but they were 12% distant from a Chinese specimen (Online Resource 1, Fig. S7);

*Neuroleon egenus*: Specimens from Portugal exhibited a 14.4% p-distance from sequences from Pakistan (Online Resource 1, Fig. S8);

*Neuroleon nemausiensis*: The 11 sequences did not cluster together, with small groups aligning with other nominal species. The maximum intraspecific distance within a species-specific dendrogram reached 16.3% (at Online Resource 1, Fig. S9).

For Ascalaphinae, sequences were obtained for *Libelloides longicornis*, *L. coccajus*, and *Palpares libelluloides*. *Libelloides longicornis*: Germany specimens showed slight differences compared to those from France (0.4%), Italy (0.6%) and the Iberian Peninsula (1.4%);

*Libelloides coccajus*: Five sequences clustered with *L. longicornis* at a 0.4% distance;

*Palpares libelluloides*: three available barcodes, the specimen from Azerbaijan showed 7% distance from one Italian specimen (EU839755.1), and 9.5% from another Italian sequence (MG334607.1), identified by BD.

## 4. DISCUSSION

We detected potential KBAs for sixteen species of antlions and owlflies using 10 km x 10 km grid cells (Table 1). When applying smaller grid cells (5 km x 5 km), three species—*Myrmeleon punicanus, My. gerlindae*, and *Megistopus lucasi*—identify smaller but overlapping potential KBAs in western Sicily and Sardinia (Online Resource 1, Fig. S4). Consistent with previous findings (Farooq et al. 2023), smaller grid cells triggered fewer potential KBAs due to the reduced number of species meeting criteria. Our study further highlights limitations of commonly used methodological approaches for assessing KBAs, such as AOO and AOH mapping when applied to taxa with incomplete distributional data or specialized habitat requirements. AOO maps were fragmented, likely underrepresenting actual distributions, as they assume occurrence records comprehensively represent species’ global ranges—an assumption often invalid for understudied taxa. Similarly, AOH maps, derived from 100-meter resolution satellite data, rely on broad land-cover categories that fail to capture microhabitats critical for antlions.

The analysis of overlap between the identified potential KBAs for antlions, owlflies and the national network of PAs and Natura 2000 sites, revealed an overlap of 18.67% and 27.63%, respectively (Online Resource 1, Fig. S1 and S2). One of the purposes of KBAs is to inform decisions aimed at improving existing networks of PAs (Langhammer et al. 2007) while simultaneously identifying new sites that are relevant for the persistence of biodiversity. However, PAs often exclude relevant habitats and species from conservation planning, as they were established to safeguard charismatic vertebrate or plant species (Rodrigues et al. 2004; Pimm et al. 2014 Klein et al. 2015), leaving most insect species underrepresented (Chefaoui et al. 2005; Chowdhury et al. 2023). Our findings show that 19.94% of the identified potential KBAs overlap with the current Italian KBA map. When considering neglected groups such as antlions, new areas were identified as potentially relevant for biodiversity conservation, including those located in endangered habitats.

### 4.1 Spatial data uncertainty and potential errors

Limited attention has been given to potential errors stemming from data uncertainties, particularly for taxa with unresolved taxonomic and poor spatial information. This omission risks their exclusion from spatial-based conservation metrics. Myrmeleontidae are globally understudied, as the ecology and life history of most species are unknown, and their taxonomy remains poorly investigated. This study demonstrates how KBA methodology can be applied to insect taxa with unresolved taxonomy and fragmented distributional and molecular data— conditions representative of many insect groups (Sánchez-Fernández et al. 2021; Garcia-Rosello et al. 2023).

Our analysis revealed that all endemic and sub-endemic species triggered potential KBAs, while most species with broader distributions did not meet the criteria. Exceptions included *Neuroleon microstenus*, and *N. nemausiensis,* both widely distributed (Aspöck et al. 1980). However, these results may be biased by the uneven distribution of occurrence points, which are geographically clustered and limited in number. Our results further suggest that many species with wide distributions may hide cryptic lineages.

### 4.2 Genetic insights into antlions and owlfly diversity

Species barcoding revealed significant genetic diversity among geographically distant populations of antlions and owlflies. This diversity frequently exceeded the species delimitation threshold (p-distance 3%) when subjected to the Objective Clustering algorithm (Meier et al. 2006; Ratnasingham & Hebert 2013). Six antlion species showed high intraspecific p-distance, ranging from 4.46% to 16.29%. Sequences from South Caucasus for three species (*Neuroleon nemausiensis*, *Distoleon tetragrammicus*, and *Palpares libelluloides*) showed substantial genetic divergence from European populations (Online Resource 1, Fig. S5 and S6). Similarly, high intraspecific p-distance observed between European and Chinese populations of *Dendroleon pantherinus* likely indicates a misidentification; Chinese records of *De. pantherinus* were instead attributable to the closely related *De. similis* (Krivokhatsky 2011).

For Ascalaphinae, barcode gap analysis identified two main clusters of *Libelloides longicornis* at a p-distance of 1.41%. Furthermore, sequences of *Libelloides coccajus* nested within a *L. longicornis* cluster, suggesting unresolved taxonomy. Assuming no sequencing errors, these results imply that species with high intraspecific molecular diversity may conceal cryptic lineages or represent misidentifications. Such taxonomic uncertainties directly reflect species distribution ranges, with cascading implications for ecology and conservation studies (Bortolus 2008). Addressing these gaps requires comprehensive taxonomic revisions and increased sampling efforts (Meier & Dikow 2004; Coddington et al. 2009).

### 4.3 Conservation implications for coastal sand dunes

Coastal sand dunes deliver key ecosystem services, including shoreline protection, nutrient regulation, carbon storage, beside many recreational opportunities (Maes et al. 2012; Arcidiacono et al. 2016). However, they face significant threats from both natural and anthropogenic disturbances (Nield & Baas 2008; Crain et al. 2009; Jackson & Cooper 2011; Pranzini & Williams 2013). The impact of anthropogenic activities on insects inhabiting arid ecosystems, including coastal dunes, remains largely unexplored.

Antlions and owlflies are key representatives of these ecosystems. Many species are stenoecious, exhibiting high habitat specificity. Antlion larvae, in particular, are strongly selective microhabitat specialists, often found in sandy habitats like coastal dunes due to their burrowing behaviour. While adult antlions are capable of wide dispersal, larvae occupy well-defined niches minimizing interspecific competition (Stange & Miller 1990; Devetak 2000; Stange 2003; Barkae et al. 2012; Badano & Pantaleoni 2014a,b). Habitat-specific features, including substrate granulometry, vegetation, and exposure to sunlight, influence larval distributions and behaviors. These ecological traits make antlions valuable bioindicators for assessing the impacts of habitat alteration.

Based on our results, four dune specialists triggered potential KBAs in sites with dune habitats: *Myrmeleon mariaemathildae*, *Megistopus lucasi*, *Neuroleon ochreatus*, and *Synclisis baetica*. These species are associated with dune habitats, usually but not exclusively on the coast, typically with sparse vegetation. Among them, *Synclisis baetica* has the widest distribution, with larvae that are active predators in exposed sandy environments. Conversely, *Megistopus lucasi* is known for a handful of localities in Sardinia and in Central Italy with well-preserved dune systems, and over one century old records from North Africa. Finally, *Myrmeleon mariaemathildae* is a recently described species restricted to the Thyrrenian islands and Tunisia, while *Neuroleon ochreatus* is limited to Mediterranean Western Europe.

## 5. CONCLUSIONS

The limitations of the KBA approach highlighted in this study apply to many overlooked taxa, such as antlions and owlflies, complicating the development of comprehensive KBA network on a global scale. These challenges emphasize the need to integrate neglected taxa into conservation planning, as they provide insights into ecosystem health and biodiversity patterns. Additionally, integrating genetic data into KBA assessments, as demonstrated here, can uncover cryptic diversity and evolutionary significance, ultimately enabling more effective conservation strategies.

## STATEMENTS AND DECLARATIONS

### Funding

This work received support from The European Union–NextGenerationEU as part of the National Biodiversity Future Center, Italian National Recovery and Resilience Plan (NRRP) Mission 4 Component 2 Investment 1.4 (CUP: B83C22002950007); by the Italian Ministry of University and Research (MUR) under the PRIN Project “Identification of priority sites for the conservation of terrestrial animal and plant diversity to meet European and CBD 2030 targets” CUP B53D23011990006 and by the NBFC to University of Siena/Department of Life Sciences, funded by the Italian Ministry of University and Research, PNRR, Missione 4 Componente 2, ‘Dalla ricerca all’impresa’, Investimento 1.4, Project CN00000033.

### Competing Interests

The authors have no relevant financial or non-financial interests to disclose.

### Author contributions

All authors contributed to the study conception and design. Data curation was carried out by Aleida Ascenzi, Davide Badano, Dario Nania, and Andrea Cristiano. Aleida Ascenzi, Dario Nania, and Andrea Cristiano contributed to the investigation and methodology. Project administration and funding acquisition were managed by Pierfilippo Cerretti and Michela Pacifici. Formal analyses were performed by Andrea Cristiano. Visualization was handled by Aleida Ascenzi, Dario Nania, Andrea Cristiano, and Davide Badano. The original draft of the manuscript was written by Aleida Ascenzi and Dario Nania, with critical revisions and editing provided by all the other authors. All authors approved the final manuscript.

## Supporting information

Online Resource 1

Online Resource 2

Online Resource 3

Online Resource 4

Online Resource 5

## ACKNOWLEDGEMENTS

We acknowledge the European Union–NextGenerationEU, the Italian Ministry of University and Research (MUR). Special thanks to Roberto A. Pantaleoni (IRET CNR SS, Italy) for the support to DB in taking the photos of some larval specimens.

## Notes

### Competing Interest Statement

The authors have declared no competing interest.

