## Supplementary material for "Neglected predatory insects trigger potential Key Biodiversity Areas in threatened coastal habitats": Online Resource 1

Biodiversity and Conservation

Aleida Ascenzi*[1,2], Dario Nania[1,2], Andrea Cristiano[1,3], Davide Badano[4,5], Michela Pacifici[1], Pierfilippo Cerretti[1,2]

[1]Department of Biology and Biotechnologies, Sapienza University of Rome, Rome, Italy

[2]Museum of Zoology (MZUR), Sapienza University of Rome, Rome, Italy

[3]Department of Geography and Environmental Sciences, Northumbria University, Newcastle upon Tyne, United Kingdom

[4]Department of Life Sciences, University of Siena, Siena, Italy

[5]NBFC, National Biodiversity Future Center, Palermo, Italy

*Corresponding author

*Aleida Ascenzi: 0009-0001-3056-2966 ;

**Table S1** List of the species of Myrmeleontidae and Ascalaphidae included in the work, with details on distribution of occurrences across data sources

| Scientific_name | Occurrence points | Literature | Collected | GBIF | iNaturalist |
|---|---|---|---|---|---|
| *Creoleon_corsicus* | 31 | 25 | 0 | 4 | 2 |
| *Creoleon_lugdunensis* | 284 | 248 | 5 | 0 | 31 |
| *Deleproctophylla_australis* | 54 | 50 | 4 | 0 | 0 |
| *Dendroleon_pantherinus* | 98 | 74 | 3 | 0 | 21 |
| *Distoleon_annulatus* | 16 | 16 | 0 | 0 | 0 |
| *Distoleon_tetragrammicus* | 440 | 366 | 34 | | 40 |
| *Libelloides_coccajus* | 225 | 220 | 5 | 0 | 0 |
| *Libelloides_corsicus* | 78 | 32 | 3 | 43 | 0 |
| *Libelloides_lacteus* | 96 | 95 | 1 | 0 | 0 |

| | | | | | |
|---|---|---|---|---|---|
| *Libelloides_latinus* | 73 | 66 | 7 | 0 | 0 |
| *Libelloides_longicornis* | 144 | 139 | 5 | 0 | 0 |
| *Libelloides_siculus* | 121 | 62 | 2 | 57 | 0 |
| *Macronemurus_appendiculatus* | 409 | 317 | 19 | 0 | 73 |
| *Megistopus_flavicornis* | 155 | 136 | 3 | 0 | 16 |
| *Megistopus_lucasi* | 16 | 12 | 4 | 0 | 0 |
| *Myrmeleon_gerlindae* | 7 | 5 | 2 | 0 | 0 |
| *Myrmeleon_mariaemathildae* | 39 | 37 | 2 | 0 | 0 |
| *Myrmeleon_punicanus* | 6 | 3 | 3 | 0 | 0 |
| *Neuroleon_arenarius* | 58 | 45 | 7 | 0 | 6 |
| *Neuroleon_egenus* | 47 | 36 | 10 | 0 | 1 |
| *Neuroleon_microstenus* | 55 | 40 | 10 | 0 | 5 |
| *Neuroleon_nemausiensis* | 86 | 81 | 4 | 0 | 1 |
| *Neuroleon_ochreatus* | 36 | 21 | 14 | 0 | 1 |
| *Palpares_libelluloides* | 349 | 238 | 9 | 1 | 101 |
| *Solter_liber* | 23 | 4 | 1 | 18 | 0 |
| *Synclisis_baetica* | 144 | 96 | 19 | 0 | 29 |

**Section S2** Area of habitat maps

We produced species distribution ranges by overlapping species' occurrence points with a map of world administrative boundaries. We retained only the administrative regions overlapping with the species' occurrence points to produce the range map. The ranges were generated as an intermediate step to produce the AOH maps.

Landcover and elevation data was combined to produce a base map containing both land cover and elevation data. The resulting map was masked with the species distribution ranges, and reclassified according to the species' habitat requirements to produce the AOH maps. We performed an hypergeometric distribution test to validate the 23 AOH maps (Jiménez & Soberón 2020; Dahal et al. 2022). Such an approach is based on the probability of a species' occurrence point to fall within the estimated AOH. It is a two-steps validation: (I) a model-based evaluation of habitat prevalence (i.e, the proportion of suitable habitat within a species' range), and (II) a validation using species occurrence records (presence-only) data.

**Fig. S1** Intersection of potential KBAs for Myrmeleontidae and the Italian network of protected areas

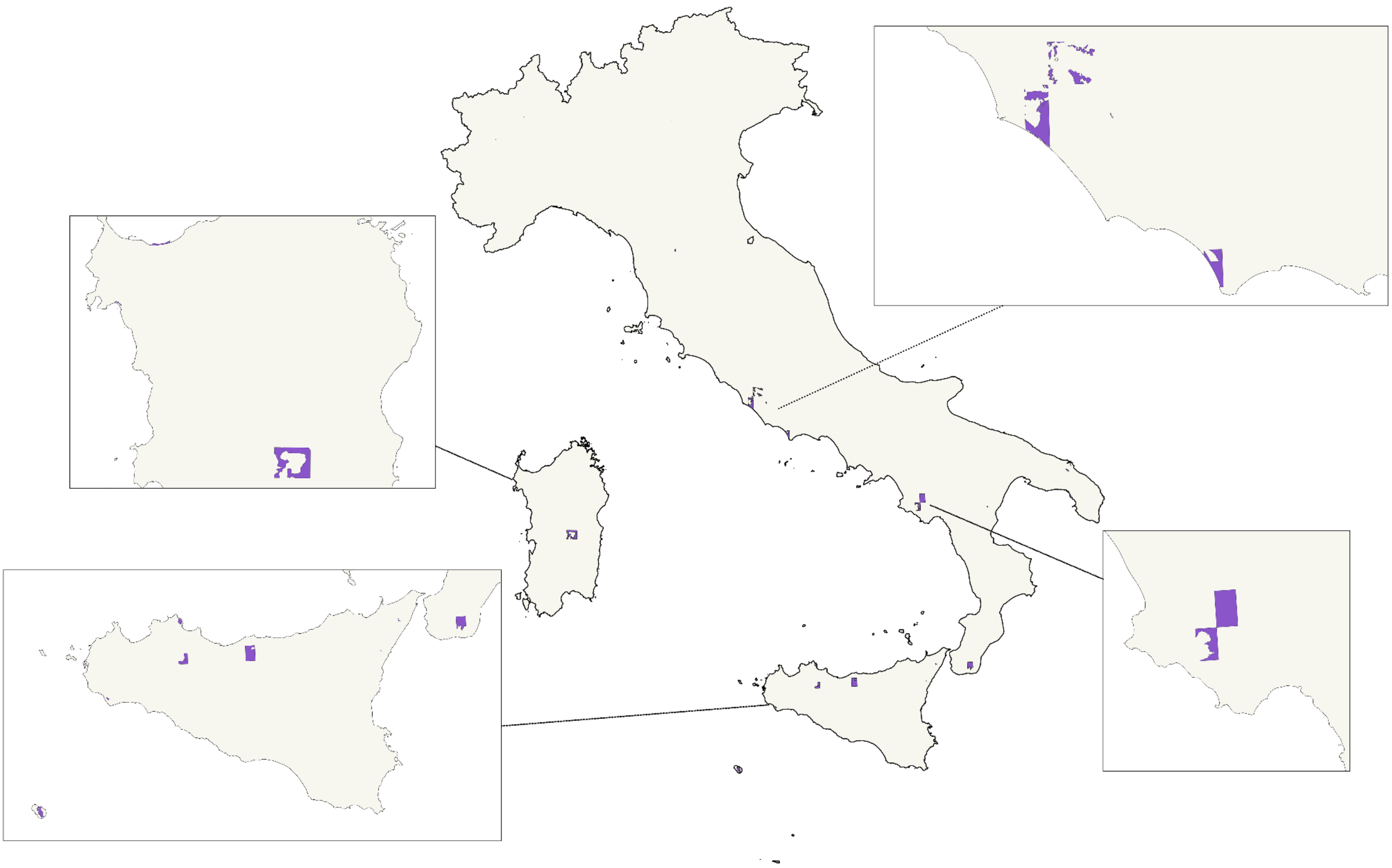

**Fig. S2** Intersection of potential KBAs for Myrmeleontidae and the Natura 2000 sites in Italy

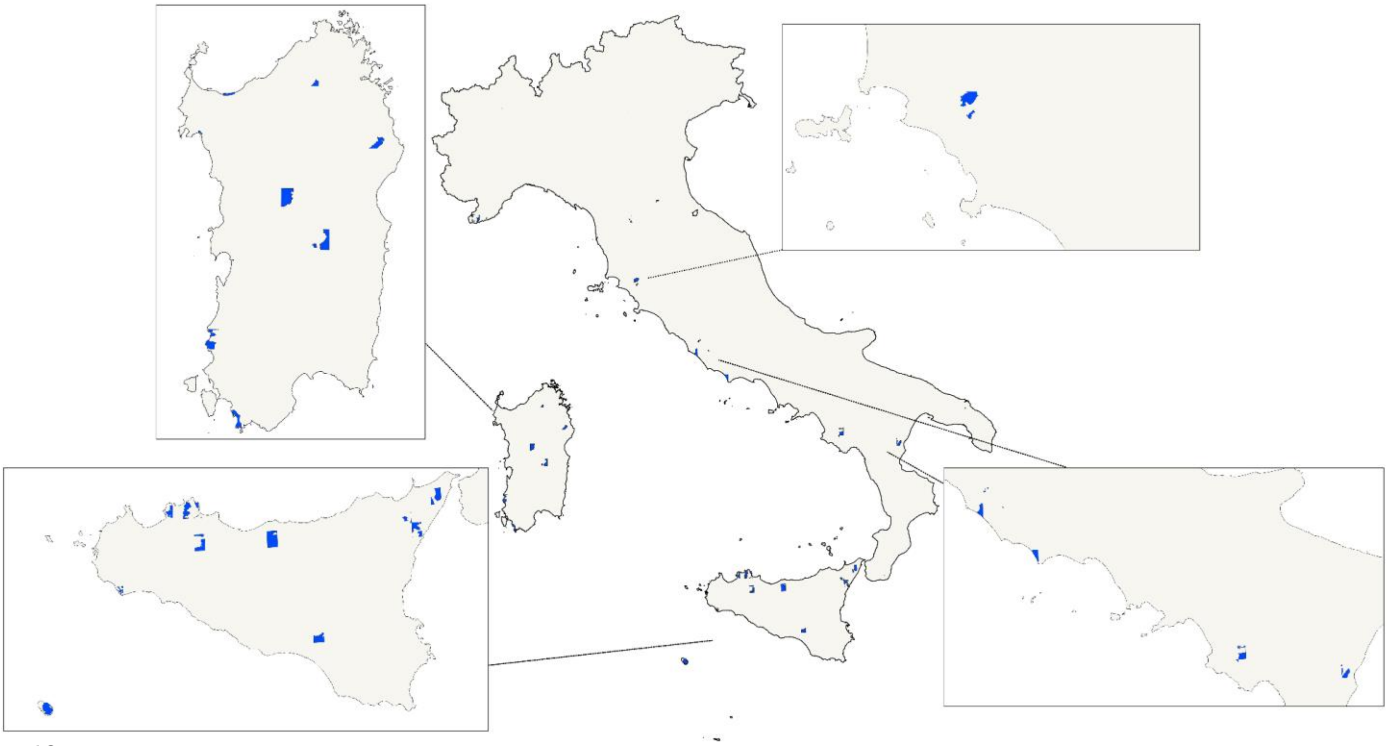

**Fig. S3** Intersection of potential KBAs for Myrmeleontidae and the current Italian KBA network

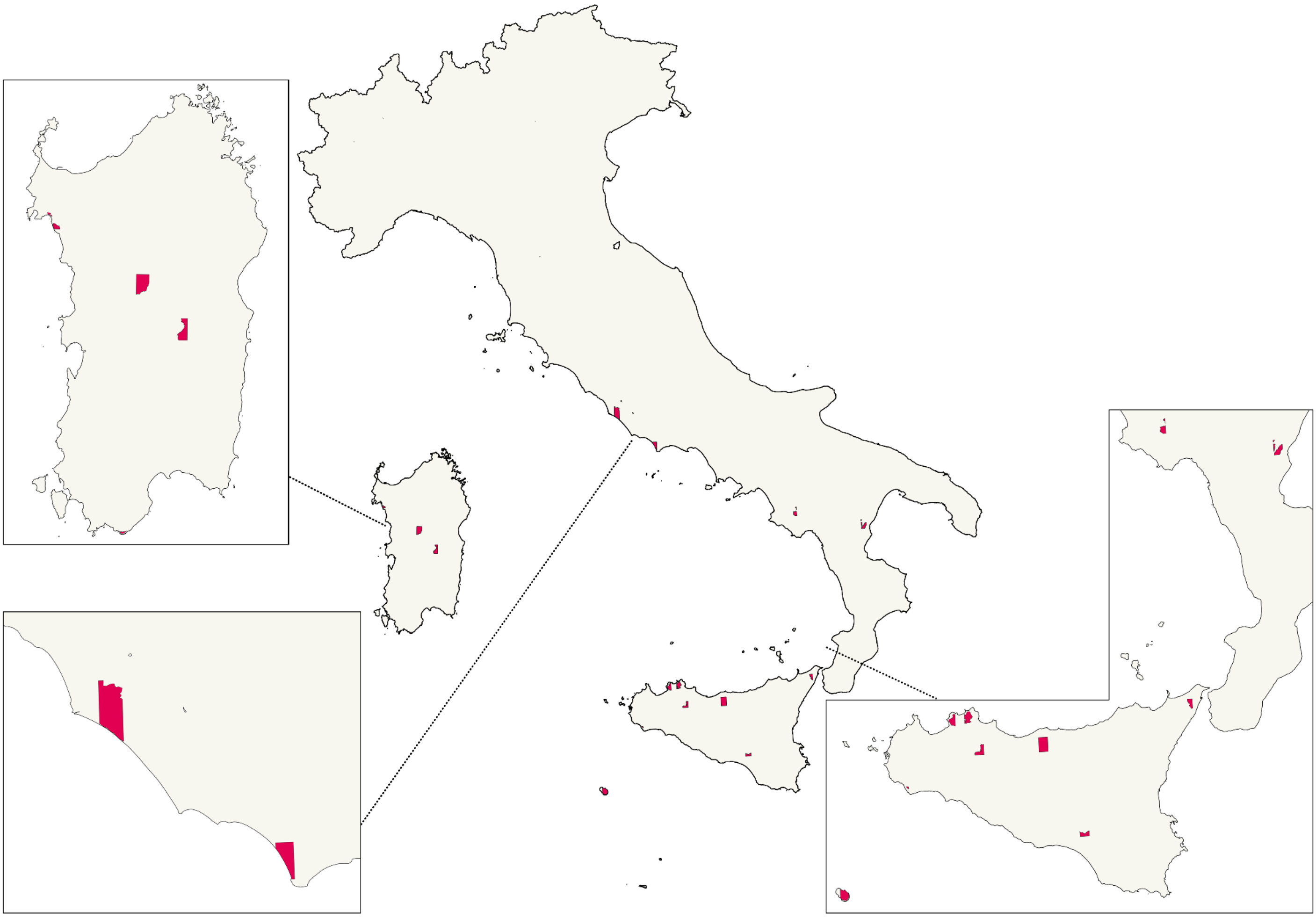

**Fig. S4.** Potential KBAs in Italy for antlions and owlflies with details on triggered criteria, generated using a 5 x 5 km cell size grid.

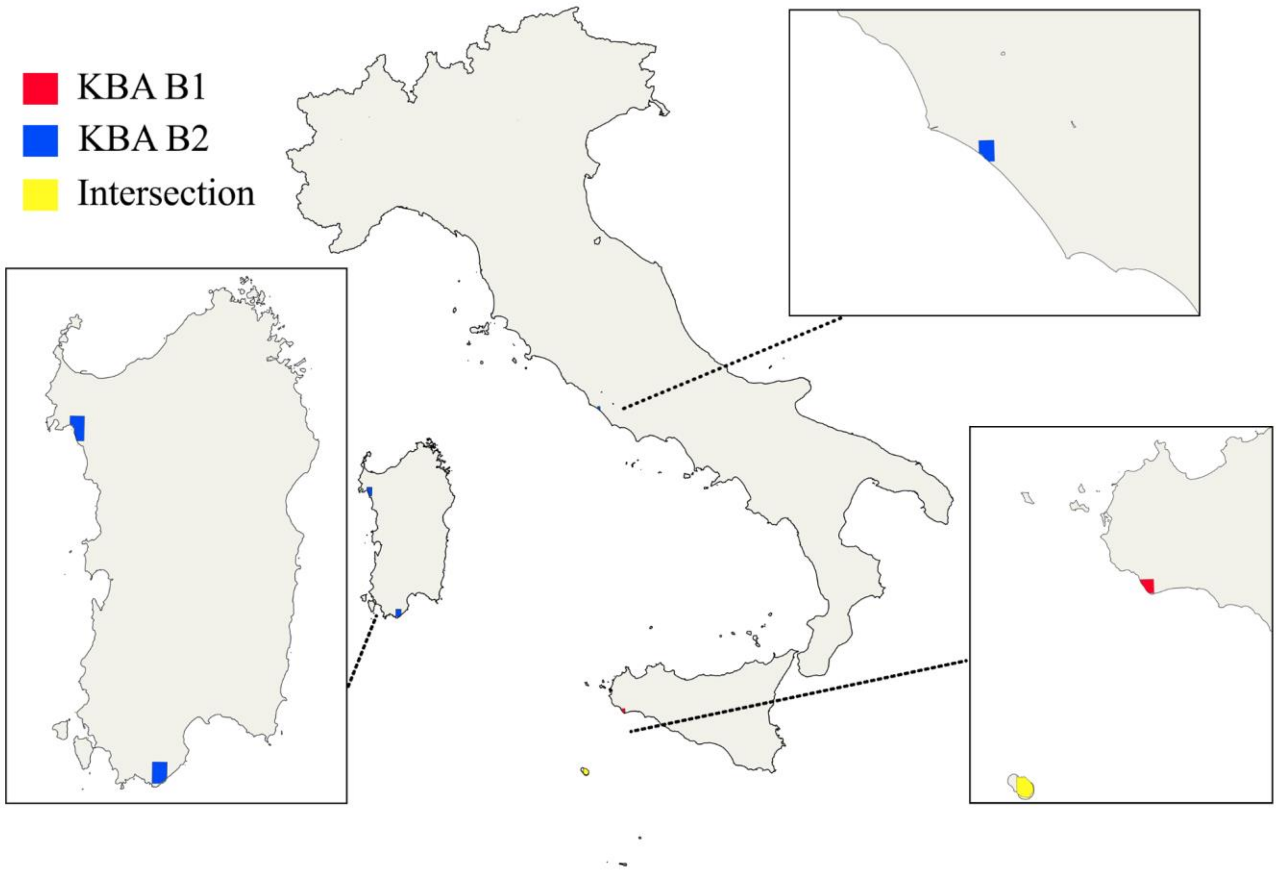

**Fig. S5.** Dendrogram of Ascalaphinae COI barcodes, constructed using Objective clustering with an uncorrected p-distance threshold of 3%. The numbers on the nodes represent the percentage difference in nucleotide bases between the clustered sequences

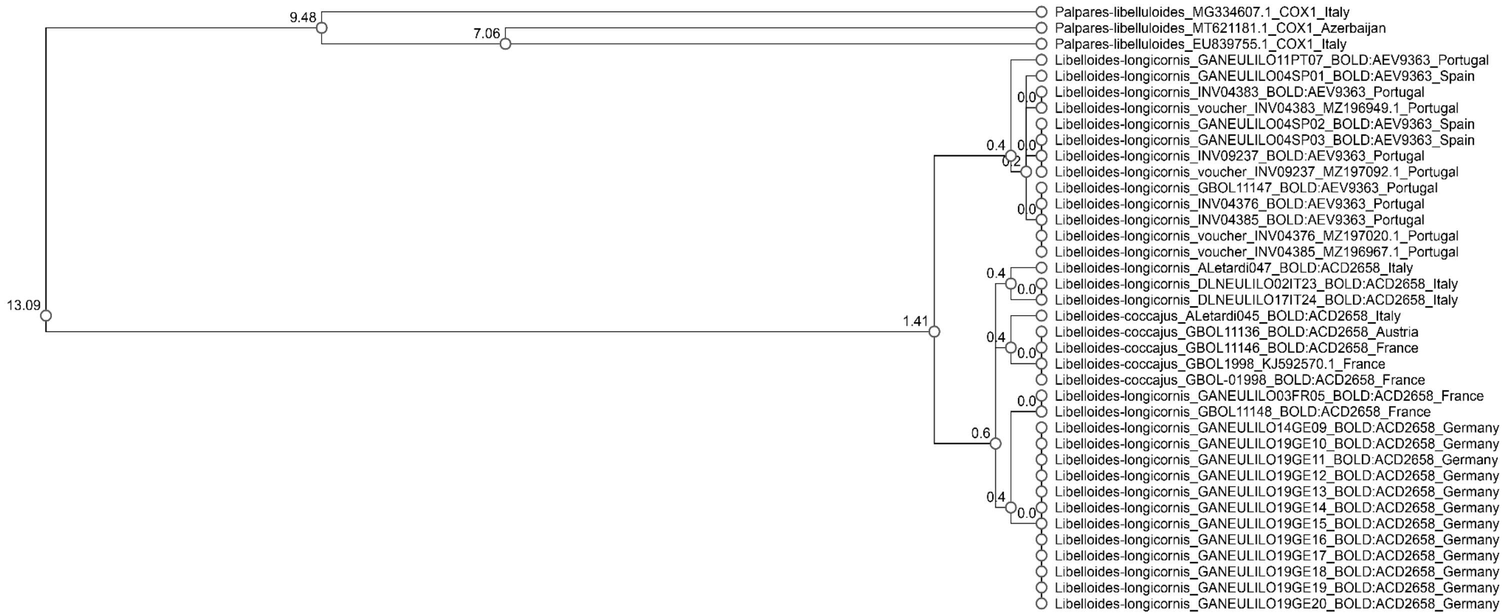

**Fig. S5** Dendrogram of Myrmeleontinae, Dendroleontinae, and Nemoleontinae COI barcodes, constructed using Objective clustering with an uncorrected p-distance threshold of 3%. The numbers on the nodes represent the percentage difference in nucleotide bases between the clustered sequences

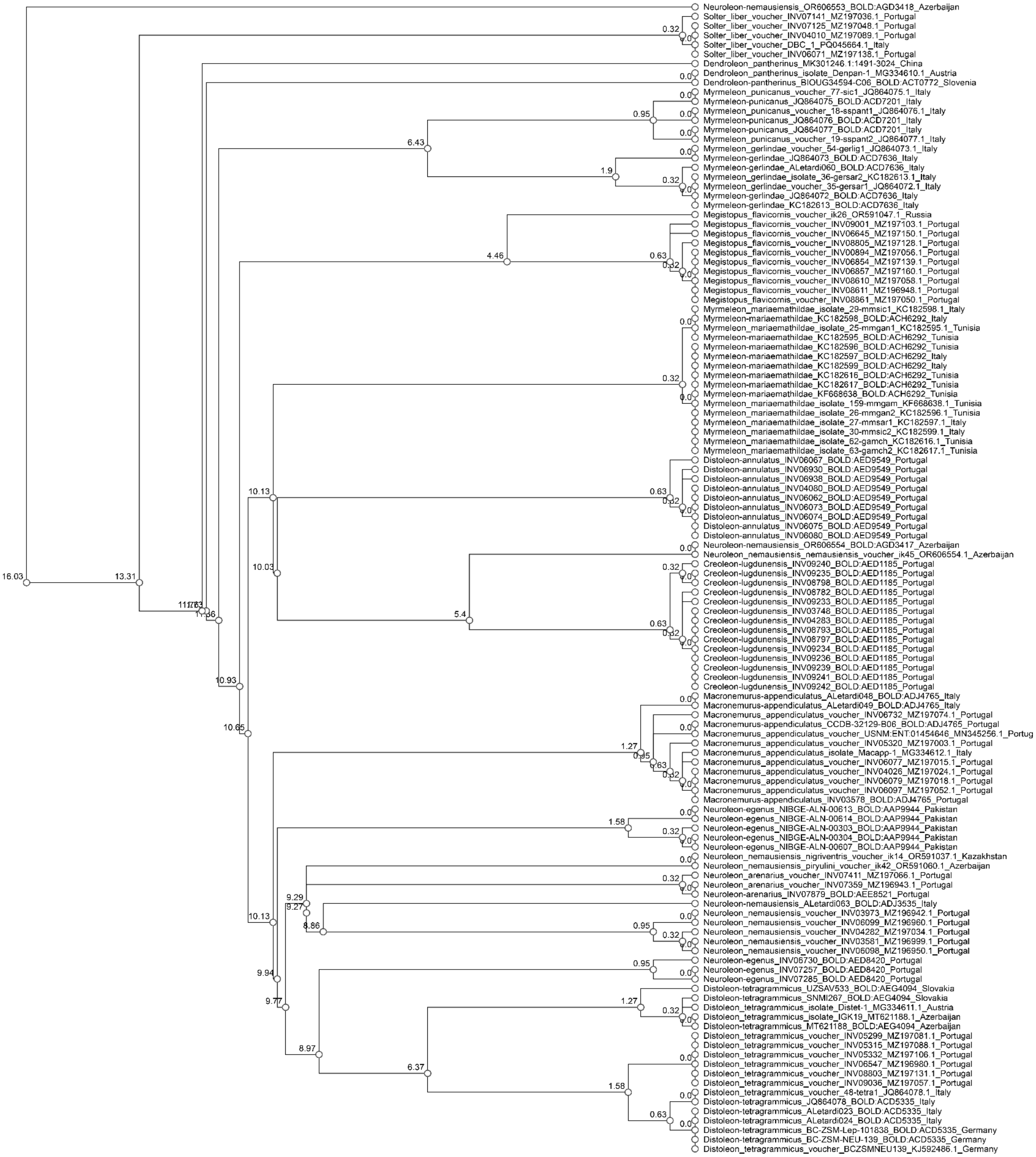

**Fig. S7** Dendrogram of *Dendroleon pantherinus* COI barcodes, constructed using Objective clustering with an uncorrected p-distance threshold of 3%. The numbers on the nodes represent the percentage difference in nucleotide bases between the clustered sequences

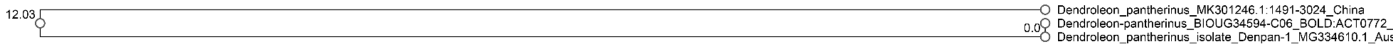


**Fig. S8** Dendrogram of *Neuroleon egenus* COI barcodes, constructed using Objective clustering with an uncorrected p-distance threshold of 3%. The numbers on the nodes represent the percentage difference in nucleotide bases between the clustered sequences

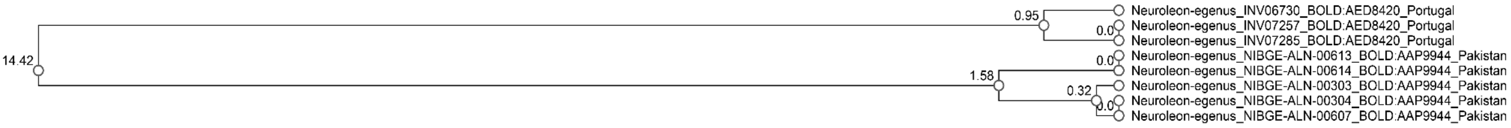


**Fig. S9** Dendrogram of *Neuroleon nemausiensis* COI barcodes, constructed using Objective clustering with an uncorrected p-distance threshold of 3%. The numbers on the nodes represent the percentage difference in nucleotide bases between the clustered sequences

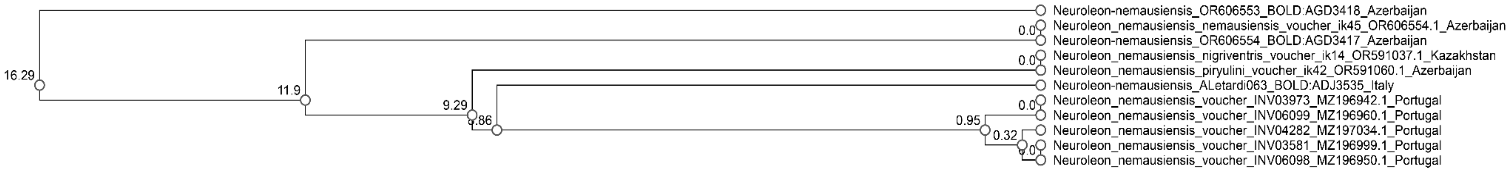


**Section S3** References list for occurrence points gathered from the literature


Acevedo Ramos F, Monserrat VJ, Contreras-Ramos A, Pérez-González S (2020) Comparative study of sensilla and other tegumentary structures of Myrmeleontidae larvae (Insecta, Neuroptera). J Morphol 281:1191-1209. https://doi.org/10.1002/jmor.21240

Aistleitner E (1980) Die Arten des Genus *Libelloides* TJEDER, 1972, der Iberischen Halbinsel (Neuroptera, Planipennia, Ascalaphidae). Taxonomie, Arealkunde, Phaenologie, Habitatwahl. (3. Beitrag zur Kenntnis der Entomofauna der Iberischen Halbinsel). Entomofauna 0001:234-297

Aistleitner E (1981) Eine neue Unterart von *Libelloides longicornis* (L.) aus den Südostalpen (Neuropteroidea, Planipennia, Ascalaphidae). Entomofauna 0002:191-202

Angelini B. (1827) Ascalafi italiani con nuova specie. Biblioteca Italiana 47:466-468

Aspöck H (1963) Über die von H. Loipetseder in Auer (Südtirol) aufgefundenen Neuropteren. Naturkundliches Jahrbuch der Stadt Linz 1963b:403-404

Aspöck H (1963) Zur Frage boreoalpiner Verbreitung bei Neuropteren. Nachrichtenblatt der Bayerischen Entomologen. 1963a:12:81–88

Aspöck H, Aspöck U, Hölzel H, Rausch H (1980) Die Neuropteren Europas: eine zusammenfassende Darstellung der Systematik, Oekologie und Chorologie der Neuropteroidea (Megaloptera, Raphidioptera, Planipennia) Europas. Goecke and Evers, Krefeld, Germany. 495 + 355 pp.

Aspöck H, Hölzel H, Aspöck U (1976) Taxonomie und Chorologie von Libelloides ictericus (Charpentier) s. l. (Neuroptera, Planipennia, Ascalaphidae) – ein Beitrag zur Biogeographie des westlichen Mittelmeerraumes. Zeitschrift der Arbeitsgemeinschaft Österreichischer Entomologen 28:17-32

Badano D (2006) Neurotteri della Liguria. Tesi di Laurea triennale in Scienze Naturali [=Undergraduate Thesis]. Università di Genova. 96 pp.

Badano D (2008) Contributo alla conoscenza dei Neurotteri della Liguria (Insecta, Neuropterida). Annali del Museo Civico di Storia Naturale Giacomo Doria 99:535-548

Badano D, Pantaleoni R (2014) The larvae of European Myrmeleontidae (Neuroptera). Zootaxa. 3762(1):1-71. https://doi.org/10.11646/zootaxa.3762.1.1

Badano D, Aspöck U, Aspöck H, Cerretti P (2017a) Phylogeny of Myrmeleontiformia based on larval morphology (Neuropterida: Neuroptera). Syst Entomol 42: 94-117. https://doi.org/10.1111/syen.12200

Badano D, Aspöck H, Aspöck U (2017b). Taxonomy and phylogeny of the genera *Gymnocnemia* Schneider, 1845, and *Megistopus* Rambur, 1842, with remarks on the systematization of the tribe Nemoleontini (Neuroptera, Myrmeleontidae). DEZ 64(1): 43-60. https://doi.org/10.3897/dez.64.11704

Brauer F (1876) Die Neuropteren Europas und insbesondere Oesterreichs mit Rücksicht auf ihre geographische Verbreitung. Pp. [263]–300. In: Festschrift zur Feier des fünfundzwanzigjährigen Bestehen Kaiserlich-Koniglichen Zoologisch-Botanischen Gesellschaf

Capra F (1934) Su alcuni odonati e mirmeleonidi di Sicilia. Ann Mus civ sor nat Giacomo Doria 57:92-97

Capra F (1937) Raccolte entomologiche nell'isola di Capraia fatte da C. Mancini e F. Capra (1927-1931). V. Odonati e Neurotteri con note sulla memoria di B. Angelini: Ascalafi italiani. Boll Soc entomol ital 69:50-58

Capra F (1976) Quanto si sa sugli Odonati e Neurotteri dell'Arcipelago Toscano (Studi sulla Riserva Naturale dell'Isola di Montecristo. XIII). Lavori della Società Italiana di Biogeografia (N.S.) 5:541-560

Carpaneto GM, Vigna Taglianti A (1995) Le attuali conoscenze sugli artropodi dei Monti Lucretili e il loro significato eco-biogeografico. De Angelis, G. (editor). Monti Lucretili Parco Regionale naturale. Invito alla lettura del territorio. 5th edition

Castellani O (1942) Neurotteri. pp. 181–183. In: Prima esplorazione entomologica del Parco Nazionale del Circeo. Inst Naz Biol Cons Naz delle Richerche, Salerno

Castellani O (1957) Contributo alla conoscenza della fauna entomologica d'italia. Neuroptera. Bollettino dell'Associazione Romana di Entomologia 11/12:9-14

Cecconi G (1908) Contributo alla fauna delle Isole Tremiti. Bollettino dei Musei di Zoologia e di Anatomia Comparata della R. Università di Torino 23(583):1-53.

Costa A (1860-1870) Fauna del Regno di Napoli ossia enumerazione di tutti gli animali che abitano le diverse regioni di questo regno e le acque che le bagnano e descrizione de' nuovi o poco esattamente conosciuti con figure ricavate da originali viventi

Costa A (1863) Nuovi studii sulla entomologia della Calabria ulteriore. Atti della Accademia delle Scienze Fisiche e Matematiche di Napoli (1)1(2):1-80

Costa A (1881) Relazione di un Viaggio nelle Calabrie per Ricerche zoologiche fatto nella state del 1876. Atti della Accademia delle Scienze Fisiche e Matematiche di Napoli (1)9(6):1-[63]

Costa A (1883) Notizie ed osservazioni sulla geo-fauna Sarda. Memoria seconda [II]. Risultamento di ricerche fatte in Sardegna nella primavera del 1882. Atti della Accademia delle Scienze Fisiche e Matematiche di Napoli (2)1(2):1-109

Costa G (1874) Fauna Salentia. Tip Ed Salentina 624 pp

Curto GM, Pantaleoni RA (1987) Note sui Myrmeleonidae (Neur. Planipennia) raccolti In oliveti del Salento (Italia meridionale). Bollettino dell’Istituto di Entomologia «Guido Grandi» dell’Università di Bologna, 42: 205-217

D'Antonio C, Fimiani P (1988) Approccio ad un inventario entomofaunistico dell'isola di Vivara (Na). Nota preliminare. (I contributo). Annuario dell'Istituto e Museo di Zoologia dell'Università di Napoli 26:155-170

De Carlini A (1885) Artropodi dell'Isola di S. Pietro. Boll Soc entomol ital 17:192-196

Dei A (1862) Insecta Senarum in agro suburbano atque in ipsa civitate lecta et in Museo Fisiocriticorum Academiae servata. Moschini, Senis. 16 pp

Ferrari G (1864) Kleiner Beitrag zur Käferfauna Venedigs und des Lido, mitgetheilt von einem süddeutschen Entomologen. Wiener Entomologische Monatsschrift 8:105-113

Ghiliani V (1874) Notizie di escursioni e di caccie entomologiche. Boll Soc entomol ital 6:91-99

Grandi G (1955) Campagna di ricerche dell'Istituto di Entomologia dell'Università di Bologna alla "Foresta Umbra" (Gargano). Annali, Accademia Italiana di Scienze Forestali 4:405-418

Griffini A (1896) Notes sur le faune entomologique piemontaise. IX. Fourmilions et Ascalaphes du Piémont. Miscellanea Entomologica 4:44-45

Güsten R (1998) Segnalazioni faunistiche. Bollettino della Società Entomologica Italiana 130(1): 80-82

Hagen HA (1860) Neuroptera Neapolitana von A. Costa, nebst Synopsis der Ascalaphen Europas. Stettiner Entomologische Zeitung 21:38-56

Hellrigl K, Hölzel H (1996) Neuropteroidea – Netzflügler. pp. 515–521. In: Hellrigl, K. (editor). Die Tierewelt Südtirols. Kommentiertes systematisch-faunistisches Verzeichnis der auf dem Gebiet der Provinz Bozen-Südtirol (Italien) bekannten Tierarten

Insom E, del Centina P, Carfì S (1985) II. Contributo alla conoscenza della fauna neurotterologica Italiana (Neuroptera: Raphidioptera, Planipennia). Redia 68:105-121

Insom E, del Centina P, Carfì S (1986) Contributo alla conoscenza della fauna neurotterologica Italiana. III. Neurotteri Planipenni della Sardegna. Redia 69:651-655

Insom E, del Centina P, Carfì S (1986) *Nicarinus poecilopterus* (Stein, 1863) (Neuroptera Myrmeleontidae) nuovo per la Sicilia. Redia 69:665-667

Krivokhatsky VA, Zakharenko AV (1994) Муравьиные львы родов Euroleon Esben-Petersen, 1918 и Kirghizoleon gen.n. (Neuroptera, Myrmeleontidae) Палеарктики [=Murav'inye l'vy rodov Euroleon Esben-Petersen, 1918 i Kirghizoleon gen. nov. (Neuroptera, Myrmeleontidae) Palearktiki; =Ant-lions of the genera Euroleon Esben-Petersen, 1918 and Kirghizoleon gen. N. (Neuroptera, Myrmeleontidae) of Palaearctic]. Энтомологическое Обозрение [=Entomologicheskoe Obozrenie; =Entomological Review, Leningrad] 73:690-699

Lacroix JL (1925) Quelques insectes Névroptères (sensu lato) recuellis en Italie. Boll Soc entomol ital 57:125-126

Lazzarini A (1896) Catalogo di Ortotteri e Neurotteri del Friuli. Pastorizia del Veneto 20-23:1-30

Letardi A (1991) Ascalafidi europei e del Medio Oriente della collezione del Museo di Zoologia dell'Università di Roma (Planipennia, Ascalaphidae). Fragmenta Entomologica, Roma 23:35-44

Letardi A, Pantaleoni RA (1996) I neuropteroidei W-Paleartici della collezione del Museo di Zoologia dell'Università di Roma (Neuropteroidea). Fragmenta Entomologica, Roma 28:277-305

Letardi A (1998) Neurotterofauna del Parco Nazionale del Cilento e Vallo di Diano (Insecta Neuropterida). Atti XVIII Congr Naz It Entomol, Maratea, 21–26 giugno 1998: 58

Letardi A (2000) [Reperti] Neuroptera, Sisyridae, Hemerobiidae, Myrmeleontidae, Ascalaphidae. Bollettino dell'Associazione Romana di Entomologia 55:144-145

Letardi A, Maltzeff P (2001) Neurotteridi e Mecotteri della Tenuta Presidenziale di Castelporziano e delle aree limitrofe (Neuroptera, Raphidioptera, Mecoptera). Bollettino dell'Associazione Romana di Entomologia 56:49-62

Letardi A (2007) Rafidiotteri, Neurotteri e Mecotteri del Parco e delle aree adiacenti (Raphidoptera, Neuroptera, Mecoptera) [=Raphidioptera, Neuroptera and Mecoptera of the park and neighbouring areas (Raphidoptera, Neuroptera, Mecoptera)]. pp. 297–305. In: Nardi, G.; Vomero, V. (editors). Artropodi del Parco Nazionale del Vesuvio: ricerche preliminari. No. 4. Conservazione Habitat Invertebrati. Cierre edizioni, Verona

Letardi A, Nicoli Aldini R (2007) Contributo alla conoscenza dei Neuropterida del Molise (Raphidioptera, Megaloptera, Neuroptera). Bollettino dell'Associazione Romana di Entomologia 62:111-130

Letardi A (2018) Contributo alla conoscenza dei Neurotteroidei italiani attraversoeventi Bioblitz (Insecta: Raphidioptera, Neuroptera). Onychium 14:99-104

Letardi A, Scalercio S (2018) Nuovi dati su Neuropterida della Calabria (Megaloptera, Raphidioptera, Neuroptera). Rivista del Museo Civico di Scienze Naturali "E. Caffi" 31:71-88

Letardi A (2022) Checklists of Italian Neuropterida (Insecta) through the last three decades. Revista Brasileira de Entomologia [=Brazilian Journal of Entomology] 66(spe):1-8 [e20220058]

Lo Valvo F (1994) Nuovi dati sulla distribuzione delle specie siciliane di Mirmeleontidae e Ascalaphidae (Insecta, Planipennia). Naturalista Siciliano (4)18:255-264

Loru L, Letardi A, Pantaleoni RA (2011) Neuropterida dell'Iglesiente (Sardegna sudoccidentale) (Raphidioptera, Neuroptera). in Nardi G; et al. (eds.). Biodiversity of Marganai and Montimannu (Sardinia). Research in the framework of the ICP Forests network. Conservazione Habitat Invertebrati 5:759-766

Marcuzzi G, Dalla Venezia L, Lorenzoni AM (1971) Appunti ecologico-qualitativi sul popolamento animale di alcuni biotopi litorali dell'alto adriatico. Atti del Reale Istituto Veneto di Scienze, Lettere ed Arti 129:119-207

Massalongo O (1891) Prospetto ragionato degli insetti della provincia di Verona. Memorie della Accademia di Verona (3)67:199-583

Massalongo O (1896) Nuova contribuzione alla fauna entomologica del veronese. Memorie della Accademia di Verona (3)72(3):47-258

Minà-Palumbo F (1858) Escursione entomologica nelle Madonie. Palingenesi 1:112

Minelli A, Mannucci MP (1993) Ricerche zoologiche nell'alto trevigiano. I. Faunistica e sinecologia di alcune cenosi dei laghi di Revine. Lavori della Societa Veneziana di Scienze Naturali 4:48-60.

Molinu A, Sassu A, Pantaleoni RA (2005) Neuropterida of the Asinara Island (NW Sardinia, Italy). Pantaleoni RA, Letardi A, Corazza C (editors). Proceedings of the Ninth International Symposium on Neuropterology. Meeting: 20-23 June 2005, Ferrara, Italy. Annali del Museo Civico di Storia Naturale di Ferrara 8:111-115

Monserrat VJ (1980) Contribución al conocimiento de los Neurópteros de Italia (Neuroptera, Planipennia). Neuroptera International 1:48-64

Navás L (1906) Notas entomológicas. XIII. Sobre una pequeña colección de Neurópteros de Italia, reunida por el profesor Felipe Silvestri, de Portici. Boletín de la [Real] Sociedad Española de Historia Natural 6:101-102

Navás L (1913-1915) Materiali per una fauna dell'arcipelago Toscano. IX. Algunos Neurópteros de la Isla del Giglio. Annali del Museo Civico di Storia Naturale Giacomo Doria 46:276-278

Navás L (1928) Insetti europei del Museo Civico di Genova. Bollettino della Società Entomologica Italiana 60:75-83

Navás L (1932) Missione scientifica del Prof. E. Zavattari nel Fezzan (1931). Neurotteri e affini. Boll Soc entomol ital 64:110-114

Navás L (1933) Insetti Neurotteri e affini del Piemonte. Memorie della Società Entomologica Italiana 12:150-162

Navás L (1935) Insectos del Piamonte (Italia). Neurópteros y afines. Boletín de la Sociedad Ibérica de Ciencias Naturales 34:33-45

Nicoli Aldini R (1983) Note sulla geonemia di alcuni Neurotteri Planipenni italiani. Giornale Italiano di Entomologia 1:123-127

Nicoli Aldini R (1992) L' “organo di Eltringham” e le correlate strutture del I urotergo in Euroleon nostras (Fourcroy) (Neuroptera Planipennia, Myrmeleontidae), al microscopio elettronico a scansione. Bollettino di Zoologia Agraria e di Bachicoltura (2)24:23-35

Nicoli Aldini R (1994) Ricerche faunistiche sui Neurotteri Planipenni di Val Camonica (Alpi e Prealpi Lombarde). Pp. 211–216. In: Atti XVII Congresso Nazionale Italiano di Entomologia. Meeting: 13-18 June 1994, Udine, Italy

Nicoli Aldini R, Baviera C (2001) Nuove acquisizioni sulla neurotterofauna di Sicilia e isole circumsiciliane (Insecta Neuropterida) [=New or interesting reports of Insecta Neuropterida in Sicily and surrounding islands]. Naturalista Siciliano (4)25:345-376

Nicoli Aldini R (2005) I neurotteri (Insecta Neuropterida) della Val Camonica e del Parco Naturale dell'Adamello (Alpi e Prealpi Lombarde): materiali per un'analisi zoogeografica del popolamento. Biogeographia, Lavori della Società Italiana di Biogeografia 26:417-430

Pantaleoni RA (1982) Neuroptera Planipennia del comprensorio delle Valli di Comacchio: indagine ecologica. Bollettino dell'Istituto di Entomologia dell'Università di Bologna Vol. XXXVII. 1-73

Pantaleoni RA (1984) Note su alcuni parassiti (s.l.) di Neurotteri Planipenni con segnalazione del ritrovamento di Acari foretici su di un Crisopide. Bollettino dell'Istituto di Entomologia "Guido Grandi" della Università degli studi di Bologna 38:193-203

Pantaleoni RA, Campadelli G, Crudele G (1994) Nuovi dati sui Neurotteri dell'alto Appennino romagnolo [=New data on the Neuropteroidea in the high Apennines of Romagna]. Bollettino dell'Istituto di Entomologia "Guido Grandi" della Università degli studi di Bologna 48:171-183

Pantaleoni RA (1986) Neurotteri dell'Italia meridionale ed insulare. Animalia, Catania 13:167-183

Pantaleoni RA (1989) Indagini ecologiche sulle popolazioni di Neurotteri (Neuropteroidea) della valle del Bidente-Ronco (Appennino Romagnolo). Doctoral dissertation. Fac Agraria, Univ Bologna.

Pantaleoni RA (1990a) I Neurotteri (Neuropteroidea) della Valle del Bidente-Ronco (Appennino Romagnolo) [=Neuropteroidea of the Bidente-Ronco Valley (Apennine of Romagna)]. Bollettino dell'Istituto di Entomologia "Guido Grandi" della Università degli studi di Bologna 44:89-142

Pantaleoni RA (1990b). I neurotteri (Insecta Neuropteroidea) delle collezioni "Zangheri" (Museo di Storia Naturale della Romagna) e "Malmerendi" (Museo civico di Scienze Naturali di Faenza). Bollettino del Museo Civico di Storia Naturale di Verona 17:277-291

Pantaleoni RA, Curto GG (1990) [Segnalazioni faunistiche italiane.] 153 - *Myrmeleon (Morter) inconspicuus* Rambur, 1842 (Planipennia Myrmeleonidae). Bollettino della Società Entomologica Italiana 121:232

Pantaleoni RA, Lo Valvo F (1995) Arthropoda di Lampedusa, Linosa e Pantelleria (Canale di Sicilia, Mar Mediterraneo). Neuroptera [=Neuroptera from Pantelleria, Lampedusa and Linosa Islands. Neuroptera]. Naturalista Siciliano (4)19(suppl.):351-356

Pantaleoni RA, Letardi A (1997) Neuropteroidea. Pp. 235–237. In: Zapparoli, M. (editor). Gli Insetti di Roma. Comune di Roma. Dipartimento Politiche della Qualità Ambientale. Area Risorsa Suolo e Tutela Ambiente. Quaderni dell'Ambiente. No. 6. Fratelli Palombi Editori, Roma. 360 pp

Pantaleoni RA, Letardi A (1998) I Neuropterida della collezione dell'Istituto di Entomologia "Guido Grandi" di Bologna. Bollettino dell'Istituto di Entomologia "Guido Grandi" della Università degli studi di Bologna 52:15-45

Pantaleoni RA, Cesaroni C, Nicoli Aldini R (2010) *Myrmeleon mariaemathildae* n. sp.: a new Mediterranean pit-building antlion (Neuropterida Myrmeleontidae). Bulletin of Insectology 63:91-98

Pantaleoni RA, Badano D (2012) *Myrmeleon punicanus* n.sp., a new pit-building antlion (Neuroptera Myrmeleontidae) from Sicily and Pantelleria. BULLETIN OF INSECTOLOGY, 65(1), 139-148

Poggi R (1993) Catalogo dei tipi di Neurotteroidei del Museo Civico di Storia Naturale "G. Doria" di Genova (Insecta) [=Catalogue of the Neuropteroidea types in the Museo Civico di Storia Naturale "G. Doria", Genoa (Insecta)]. Annali del Museo Civico di Storia Naturale Giacomo Doria 89:415-444

Pollini C (1816) Viaggio al Lago di Garda e al Monte Balbo in cui si ragiona delle cose naturali di quei luoghi aggiuntovi un cenno sulle curiosità del Bolca e degli altri monti veronesi. Mainardi, Verona. 152 pp.

Principi MM (1943) Contributi allo studio dei Neurotteri italiani. II. *Myrmeleon inconspicuus* Ramb. e Euroleon nostras Fourcroy - Boll Ist Entomol, Univ Bologna 14: pp. 131-192

Principi MM (1947) Contributi allo studio dei Neurotteri Italiani. VI. *Synclisis baetica* Ramb. (Myrmeleonidae - Acanthaclisini). Bollettino dell'Istituto di Entomologia della Università degli Studi di Bologna 16:234-253

Principi MM (1952) Ricerche zoologiche sul Massiccio del Pollino (Lucania-Calabria). VI. Neurotteri. Annuario dell'Istituto e Museo di Zoologia dell'Università di Napoli 4(10):1-22

Principi MM (1958a) Neurotteri dei Monti Sibillini (Appennino umbro-marchigiano). Memorie del Museo Civico di Storia Naturale di Verona 6:175-189

Principi MM (1958b) Il volo degli Ascalafidi. Natura e Montagna 5:54-56

Principi MM (1966) Contributi allo studio dei Neurotteri Italiani. XVIII. Neurotteri della Basilicata, della Calabria e della Sicilia. Memorie del Museo Civico di Storia Naturale di Verona 14:363-388

Principi MM, Capra F (1960) Neuroptera, Chrysopidae. in Zavattari, E. (ed.) Biogeografia delle isole Pelagie. Fauna: Invertebrati. Rendiconti, Accademia Nazionale dei LX (4)11:341-342

Ramme W (1911) Entomologische Ergebnisse einer Reise nach Oberitalien und Sütirol (1910). (Neuroptera, Odonata, Orthoptera, Lepidoptera, Diptera, Hymenoptera, Rhynchota). Berliner Entomologische Zeitschrift 56:11-32

Riggio G (1886) Materiali per una fauna entomologica dell'isola di Ustica. Prima [I] contribuzione. Naturalista Siciliano 5:52-56

Rizzotti Vlach M, Zanini D (1997). Interessanti reperti entomologici nell'area baldense meridionale. Natura Alpina 48:45-56

Schmid H (1972) Neuropteren aus dem Aspromonte, zugleich ein Beitrag zur tiergeographischen Stellung dieses Gebirges. Entomologische Zeitschrift, Frankfurt am Main 82:249-253

Schneider WG (1845) Verzeichniss der von Herrn Oberlehrer Zeller im jahre 1844 in Sicilien und Italien gesammelten Neuroptera, mit beschreibung einiger neuen arten. Stettiner Entomologische Zeitung 6:338-346

Sforzi A, Angiolini C, Bacaro G, Baini M, Bartolommei P, Bastianini M, Benocci A, Biagini G, Bonari G, et al. (2019) BIOBLITZ 2013-2014 OASI DI SAN FELICE. Atti del Museo di Storia Naturale della Maremma, 24: 11-43 https://doi.org/10.5281/zenodo.4773171

Tagliapietra V, Zanocco (1998) Il progetto Bioitaly in Trentino: invertebrati. Report, Centro di Ecologia Alpina 14:1-83

Tröger EJ (1988) Weitere Nachweise von Acanthaclisis baetica Rambur (Neuropteroidea, Planipennia, Myrmeleonidae) und ihre larven in westlichen Mittelmeergebiet und in Nordwestafrika. Entomologische Zeitschrift mit Insektenbörse 98:220-224

Tröger EJ, Rezbanyai-Reser L (1998) Fundangaben von europäischen Ameisenjungfern aus der Kollektion des Natur-Museums Luzern, sowie eine Zusammenfassung der Schweizer Arten (Neuroptera: Myrmeleontidae). Entomologische Berichte Luzern, Mitteilungen der Entomologischen Gesellschaft Luzern 40:99-108

Tröger (1999) Neue Neuropteren-Funde auf Kreta – Galathea, Berichte des Kreises Nürnberger Entomologen e.V. – 5 Supp: 8 - 12

Zàngheri P (1966) Neuroptera. Pp. 816–822. In: Repertorio sistematico e topografico della flora e fauna vivente e fossile della Romagna. Tomo II. Memorie fuori serie del Museo Civico di Storia Naturale di Verona 1:481-454
